# 3DProtDTA: the deep learning model for drug-target affinity prediction based on the residue-level protein graphs

**DOI:** 10.1101/2022.11.24.517815

**Authors:** Taras Voitsitskyi, Roman Stratiichuk, Ihor Koleiev, Leonid Popryho, Zakhar Ostrovsky, Pavel Henitsoi, Ivan Khropachev, Volodymyr Vozniak, Roman Zhytar, Diana Nechepurenko, Semen Yesylevskyy, Alan Nafiev, Serhii Starosyla

## Abstract

Accurate prediction of the drug-target affinity (DTA) *in silico* is of critical importance for modern drug discovery. Computational methods of DTA prediction, applied in the early stages of drug development, are able to speed it up and cut its cost significantly. A wide range of approaches based on machine learning was recently proposed for DTA assessment. The most promising of them are based on deep learning techniques and graph neural networks to encode molecular structures. The recent breakthrough in protein structure prediction made by AlphaFold made an unprecedented amount of proteins without experimentally defined structures accessible for computational DTA prediction. In this work, we propose a new deep learning DTA model 3DProtDTA, which utilises AlphaFold structure predictions in conjunction with the graph representation of proteins. The model is superior to its rivals on common benchmarking datasets and has a potential for further improvement.

## Introduction

Modern drug discovery remains a painfully slow and expensive process despite all recent scientific and technological advancements. It usually takes several years and the estimated cost of developing a new drug may run over a billion US dollars (Roses, 2008). More than 30% of all drugs entering phase II of clinical trials and above 58% of drugs entering phase III fail (Van Norman, 2019). It was reported that among 108 new and repurposed drugs, reported as Phase II failures between 2008 and 2010, 51% were due to insufficient efficacy (Arrowsmith, 2011). This observation highlighted the need for novel *in silico* techniques that can decrease the failure rate by filtering out compounds with low predicted efficacy in the early stages of the drug discovery pipeline. In this regard, the computational methods that assess drug-target binding affinities (DTA) are of great interest (Thafar et al., 2019) because DTA is generally considered one of the best predictors of resulting drug efficacy. Accurate prediction of the DTA is of critical importance for filtering out inefficient molecules and preventing them from reaching clinical trials, thus a multitude of computational DTA techniques have been developed in recent years.

The most accurate computational estimate of DTA could be obtained from atomistic molecular dynamics simulations (either classical, quantum or hybrid) combined with one of the modern techniques of computing the free energy of ligand binding (Pinzi and Rastelli, 2019). However, accuracy comes at the cost of very high computational demands, which makes these methods generally impractical for large-scale virtual screening.

That is why the common method of choice for estimating DTA in modern drug discovery is molecular docking, which provides a reasonable compromise between accuracy and computational efficiency (Li et al., 2019). However, it is generally believed that empirical scoring functions used in molecular docking have already approached the practical limit of accuracy, which is unlikely to be improved without introducing an additional computational burden.

In order to address these drawbacks the classical machine learning (ML) methods for determining DTA were developed. These methods do not depend on computing physical interactions between the target protein and the ligand. They are purely knowledge-based and rely on the idea that similar ligands tend to interact with similar protein targets, which are encoded into the pre-trained neural networks. Applying such models to any given ligand is blazingly fast, which allows using them for screening the numbers of compounds, which are unreachable for molecular docking or MD simulations. However, even the most successful methods from this category, such as KronRLS (Pahikkala et al., 2015) and SimBoost (He et al., 2017), were recently outperformed by more modern deep learning (DL) techniques.

Seminal DL methods of assessing DTA used string representation of the target’s amino acid sequence and a simplified linear representation of the ligands using SMILES (molecular-input line-entry system), which were subsequently encoded by 1D convolutional neural networks (CNNs) and/or long-short-term memory (LSTM) blocks (Abbasi et al., 2020; Öztürk et al., 2018). The same linear representations were shown to be efficient for DTA prediction in combination with generative adversarial networks (GANs) (Zhao et al., 2020). However, it is obvious that linear representations lead to a huge loss of information and keeping the knowledge about connectivity and 3D arrangements of both the target protein and the ligands could improve the results. Indeed, the introduction of graph neural networks (GNNs), which preserve information about connectivity and the 3D arrangement of atoms, improved the scores of the models on most of the benchmarking DTA datasets (Nguyen et al., 2021).

In contrast to the sequence-based techniques, the performance of the GNNs depends strongly on the availability and accuracy of 3D structures of target proteins. The scarcity of such structures limited the size of the training datasets and hampered the progress of GNN-based DTA models. The huge success of AlphaFold (Jumper et al., 2021) in protein structure prediction opens new opportunities for developing better DTA prediction models. Accurately predicted 3D structure of druggable protein domains that don’t have experimentally resolved structures, adds unprecedented amounts of data for training ML models and improving their performance.

In this article, we proposed a new method of constructing efficient residue-level protein graphs based on the target’s 3D structure predicted by AlphaFold and selected the best GNN architectures for this kind of data. This resulted in a new deep-learning model for predicting drug-target affinities: 3DProtDTA. When applied to common benchmark datasets our model is superior to its rivals on all evaluated metrics. The perspectives of applications and further improvements of our model are discussed.

## Materials and methods

### Datasets

We evaluated our approach on two widespread benchmark datasets for DTA prediction: Davis (Davis et al., 2011) and KIBA (Tang et al., 2014).

The Davis dataset contains the pairs of kinase proteins and their respective inhibitors with experimentally determined dissociation constant (K_d_) values. K_d_ values were transformed by equation 1 and transformed scores were used as labels for benchmarking in the same way as in the baseline approaches. There are 442 proteins and 68 ligands in this dataset.

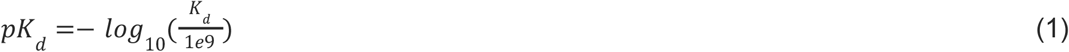

The KIBA dataset comprises scores originating from an approach called KIBA, in which inhibitor bioactivities from different sources such as K_i_, K_d_ and IC_50_ are combined. The KIBA scores were pre-processed by the SimBoost algorithm (He et al., 2017) and the final values were used as labels for model training. Initially, the KIBA dataset contained 467 proteins and 52498 ligands. For benchmarking purposes, the same authors (He et al., 2017) filtered the dataset to keep only the drugs and targets with at least 10 interactions resulting in 229 unique proteins and 2111 unique ligands.

The numbers of affinity scores and unique entries in the datasets are summarised in Table 1.

**Table 1.** Summary of the benchmark datasets.

| Dataset | Proteins | Ligands | Interactions |
| --- | --- | --- | --- |
| Davis | 442 | 68 | 30056 |
| KIBA | 229 | 2111 | 118254 |

We used isomorphic SMILES strings for both datasets and UniProt (The UniProt Consortium, 2019) accession codes for the KIBA dataset, provided by DeepDTA (Öztürk et al., 2018), as initial entries representation. For the Davis dataset, proteins with the same UniProt accession codes may represent different entries. Therefore, unique identifiers were assigned to each target entry annotated by UniProt accession code, mutation type, phosphorylation status, proteins in a complex, and protein domains. Figure 1 shows distributions of labels for Davis (a) and KIBA (b) datasets.

**Figure 1.**
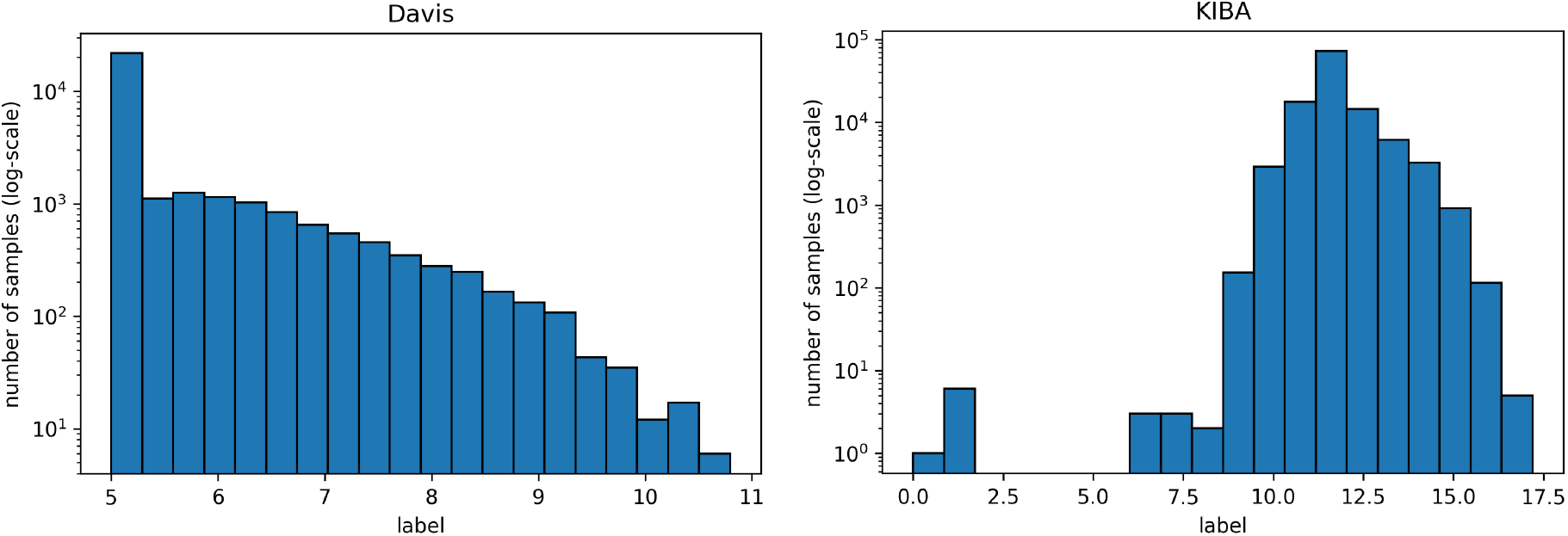
Distribution of Davis (left) and KIBA (right) labels used directly in benchmarking (note that the y-axis is log-scale).

### Ligand representation

We utilised modified molecular graphs, initially proposed in the approach for drug property prediction Chemi-Net (Liu et al., 2019) along with the standard Morgan fingerprints (Rogers and Hahn, 2010) to represent ligands for DTA prediction.

Python API of an open-source cheminformatics package RDKit v. 2021.03 was used to generate both ligand representations based on isomorphic SMILES. We calculated 1024-bit vectors of classical Morgan fingerprints with radius 2 and 1024-bit vectors of feature-based fingerprint invariants (Gobbi and Poppinger, 1998) with the same radius. Both vectors were concatenated into a single 2048-bit vector.

The graphs for ligands were generated on the atomistic level (one node in the graph is one heavy atom in a ligand). Figure 2 shows distributions of the number of heavy atoms in the used ligands, while Table 2 shows the features for a molecular graph representation of the ligands.

**Figure 2.**
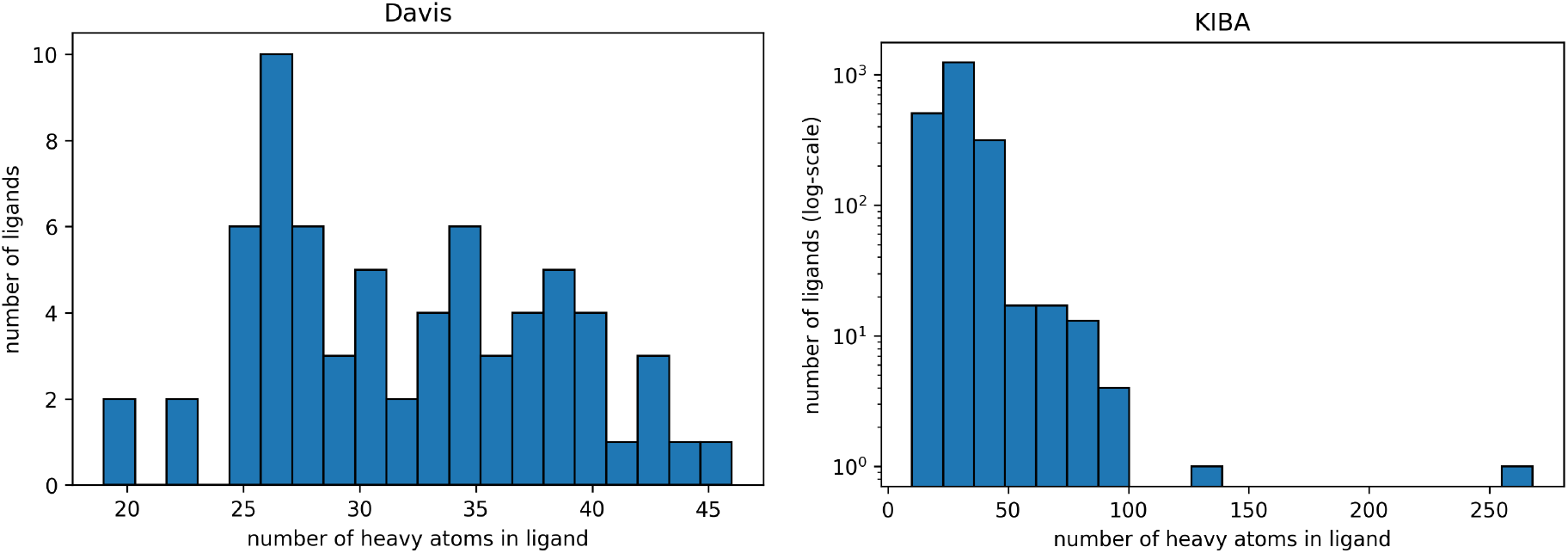
Distribution of the number of heavy atoms in the ligands in Davis (left) and KIBA (right) datasets. The Y axis is log-transformed for the KIBA dataset.

**Table 2.** Atomic-level graph features for ligand representation.

| Name | Size |
| --- | --- |
| **Node features** |  |
| One-hot encoded atom type (C, N, O, F, S, Cl, Br, P, I) | 9 |
| Atom mass (scaled by min-max) | 1 |
| Number of directly-bonded atom neighbours (scaled by min-max) | 1 |
| Total number of bonded hydrogens (scaled by min-max) | 1 |
| One-hot encoded atom hybridization (sp<sup>2</sup>, sp<sup>3</sup>) | 2 |
| Is atom in a ring (1 - yes, 0 - no) | 1 |
| Is atom aromatic (1 - yes, 0 - no) | 1 |
| Is atom hydrophobic (1 - yes, 0 - no) | 1 |
| Is atom metal (1 - yes, 0 - no) | 1 |
| Is atom halogen (1 - yes, 0 - no) | 1 |
| Is atom donor (1 - yes, 0 - no) | 1 |
| Is atom acceptor (1 - yes, 0 - no) | 1 |
| Is atom positively charged (1 - yes, 0 - no) | 1 |
| Is atom negatively charged (1 - yes, 0 - no) | 1 |
| **Edge features** |  |
| One-hot encoded bond type (single, double, triple, aromatic) | 4 |
| Bond is conjugated (1 - yes, 0 - no) | 1 |
| Bond is in the ring (1 - yes, 0 - no) | 1 |

The Morgan fingerprints and ligand graphs are available in the GitHub repository. The order of the features in the graphs is the same as in Table 2.

### Protein representation

We have developed the residue-level protein graph based on 3D protein structures generated by AlphaFold (Jumper et al., 2021). Approximately 50% of the proteins in both datasets have known 3D structures deposited in the Protein Data Bank but we decided to use AlphaFold predictions for all proteins to make our approach unified and to avoid additional tedious pre-processing of experimentally determined structures, which are often incomplete, contain irrelevant crystallographic ligands, etc.

For the KIBA dataset, all the structures were obtained from the AlphaFold protein structure database (https://alphafold.ebi.ac.uk/). For the Davis dataset only single protein entries without mutations were downloaded from the AlphaFold database. For the rest of the entries, 3D structures were generated manually using an accelerated implementation of the AlphaFold algorithm in Google Colaboratory: ColabFold (Mirdita et al., 2022).

To avoid undesirable noise from the parts of proteins, which have weak or no relation to the ligand binding, we have parsed domain annotations from UniProt (The UniProt Consortium, 2019) to determine the ligand binding sites. Both datasets contain only the kinase enzymes and the ligands with kinase inhibitor activity. Consequently, only the domains with known kinase activity or related to the kinase activity (annotated by UniProt as a protein kinase, histidine kinase, PI3K/PI4K, PIPK, AGC-kinase, or CBS) were kept in the protein structures.

This preprocessing step not only decreased the noise in the data but also eliminated most residues with a low per-residue confidence score (pLDDT). In AlphaFold a pLDDT is a continuous scale from 0 - 100 (higher is better), which shows the quality of structure prediction. pLDDT lower than 70 emerges in predicted 3D structures if they are unstructured in physiological conditions or the amino acid sequence has low alignment depth (Jumper et al., 2021). The regions with such low scores should be treated with caution. Since the domains related to kinase activity are mostly well studied and available in databases of experimental protein structures, keeping only them and removing other regions improves the average pLDDT score. Figure 3 shows the number of amino acids in processed PDB files (A) as well as the distribution of pLDDT scores (B).

**Figure 3.**
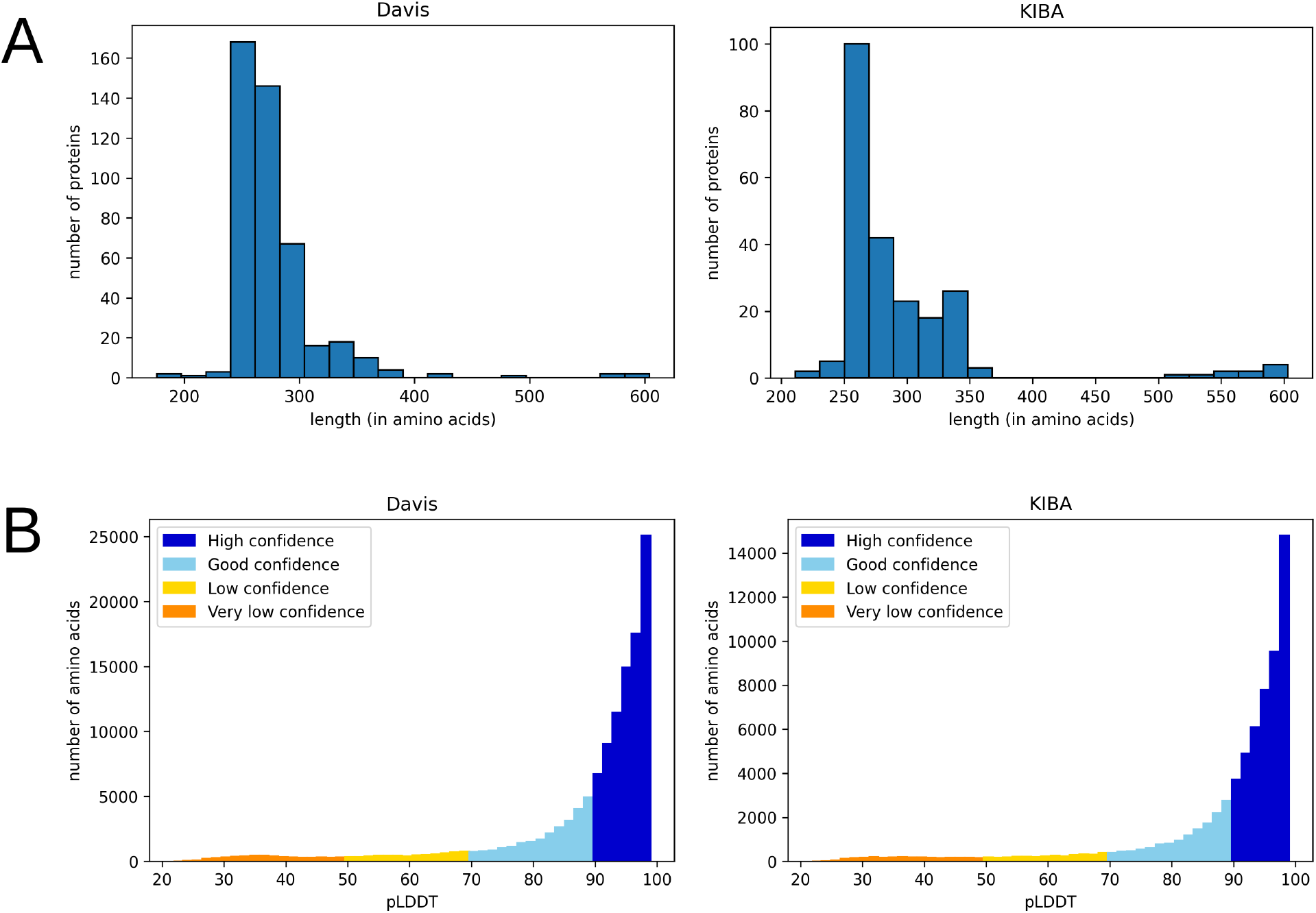
Distribution of the number of amino acids (A) and pLDDT scores (B) in processed 3D protein structures (after removal of the parts not related to kinase activity) for Davis (left) and KIBA (right) datasets.

Filtered 3D structures were converted into the residue-level graphs using Biopython v. 1.79 (Cock et al., 2009) and Pteros (Yesylevskyy, 2015, 2012). Inspired by the Open Drug Discovery Toolkit (Wójcikowski et al., 2015), we have developed an approach for encoding protein properties in the graph edge features. An edge was created if two amino acids form a either covalent bond or a non-covalent contact within a particular distance cutoff. The edge features define the type of this connectivity (Table 3). This technique allowed reducing the size of the protein graph in terms of the number of edges compared to the conventional protein graph generation approaches that define the same distance threshold for all types of residue-residue interactions.

**Table 3.** Bond types and corresponding distance cutoffs used for graph generation and assignment of edge features.

| Bond type | Distance cutoff (Å) |
| --- | --- |
| Covalent | - |
| Hydrophobic contacts | 4 |
| Hydrogen bond | 3.5 |
| Salt bridge | 4 |
| Cation-pi interaction | 5 |
| Perpendicular pi-stacking | 5 |
| Parallel pi-stacking | 5 |

Such a graph is directed because of the hydrogen bonds, salt bridges, and cation-pi interactions, which are non-commutative and require specification of the roles for each of involved residues. For example, in the edge created by the hydrogen bond between nodes *a* and *b*, it is important to identify which node is a donor and which one is an acceptor. Thus, edge a-b is assigned edge features that are different from those of edge b-a. Similarly, the salt bridges require distinguishing between cationic residue and anionic residue nodes and the cation-pi interactions - between cationic and aromatic residues. In contrast, covalent bonds, pi-stacking and hydrophobic contacts are symmetric.

For each of the 20 standard amino acid types, we assigned seven characteristics AAPHY7 (Meiler et al., 2001) and 23 blosum62 values according to alignments of homologous protein sequences (Mount, 2008) provided by GraphSol (Chen et al., 2021). In addition, the structure-dependent and sequence-dependent node and edge features were used (Table 4).

**Table 4.** Residue-level node features for protein graph representation.

| Name | Size |
| --- | --- |
| **Node features** |  |
| Solvent-accessible surface area (scaled by mean and standard deviation) | 1 |
| Phi angle (in degrees, divided by 180) | 1 |
| Psi angle (in degrees, divided by 180) | 1 |
| One-hot encoded belonging to secondary structure (alpha helix, isolated beta-bridge residue, strand, 3-10 helix, turn, bend) | 6 |
| AAPHY7 descriptors of a residue | 7 |
| blosum62 descriptors of a residue | 23 |
| Phosphorylated (1 - yes, 0 - no, same for all nodes) | 1 |
| Mutated (1 - yes, 0 - no, same for all nodes) | 1 |

| Edge features |  |
| --- | --- |
| Covalent bond (1 - yes, 0 - no) | 1 |
| Hydrophobic contact (1 - yes, 0 - no) | 1 |
| Hydrogen bond (from donor to acceptor; 1 - yes, 0 - no) | 1 |
| Hydrogen bond (from acceptor to donor; 1 - yes, 0 - no) | 1 |
| Salt bridge (from cation to anion; 1 - yes, 0 - no) | 1 |
| Salt bridge (from anion to cation; 1 - yes, 0 - no) | 1 |
| Pi-cation interaction (from aromatic ring to cation; 1 - yes, 0 - no) | 1 |
| Pi-cation interaction (from cation to aromatic ring; 1 - yes, 0 - no) | 1 |
| Parallel pi-stacking (1 - yes, 0 - no) | 1 |
| Perpendicular pi-stacking (1 - yes, 0 - no) | 1 |

As the two last node features for the protein graphs we added phosphorylation status and mutation status. Each of the two features is either 1 (phosphorylated/mutated) or 0 (non-phosphorylated/non-mutated) and is the same for each node in a protein graph. It is essential in the case of the Davis dataset due to the presence of protein entries with the same sequence but annotated as phosphorylated/non-phosphorylated or wild-type and mutant entries with internal tandem duplication (ITD) mutation that is hard to translate into sequence unambiguously. Ignoring this data would cause the situation when proteins with identical graph representation have different binding affinities to the same ligand.

Generated protein graphs are available in the GitHub repository. The order of features in the graph is the same as in Table 4.

### Model architecture

We used GNNs to extract features from the ligand and protein graphs followed by fully connected (FC) neural network layers. The general model architecture is represented in Figure 4.

**Figure 4.**
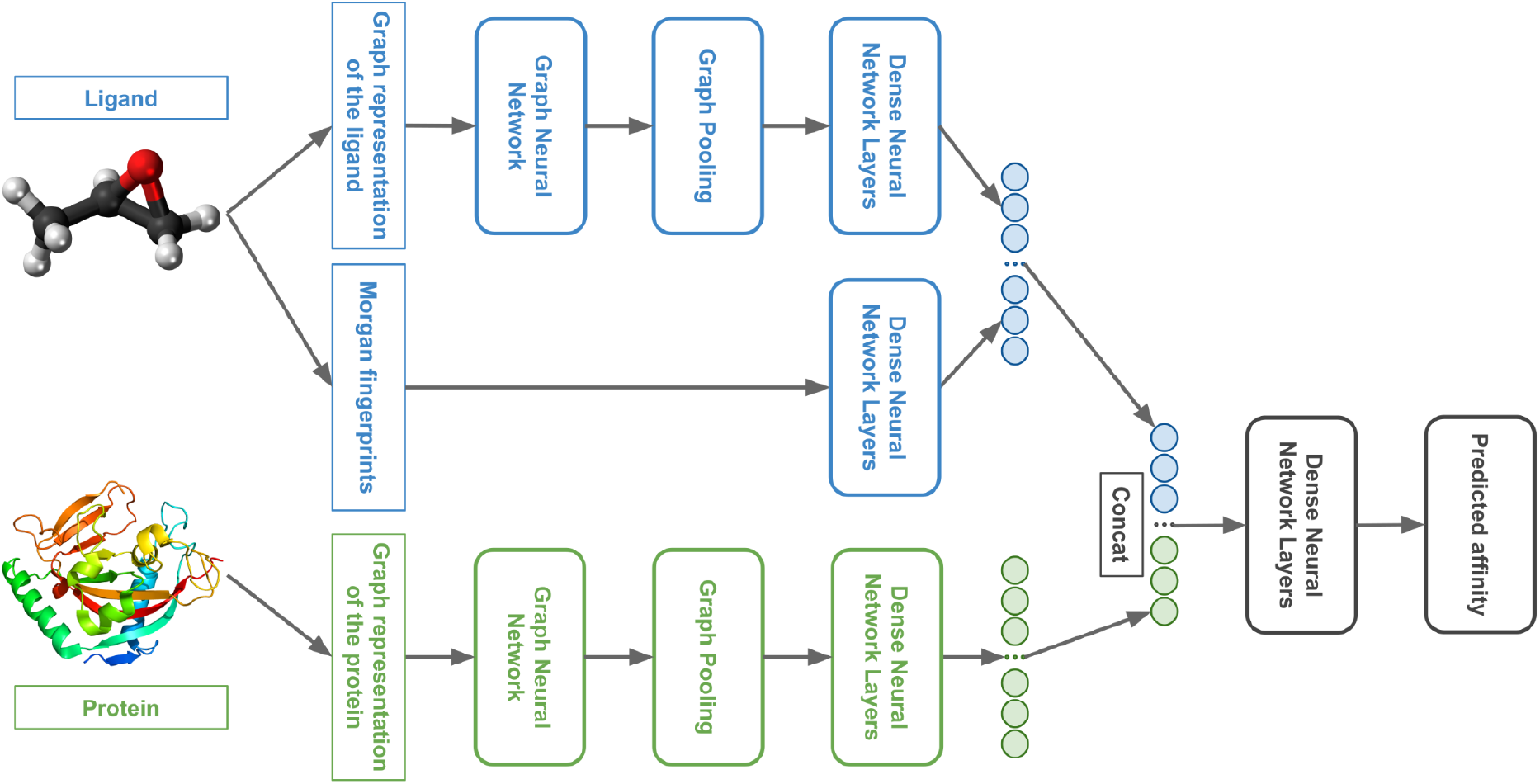
The general architecture of the 3DProtDTA model.

We tuned 3DProtDTA for the single GNN type or the combination of several GNN types that provides the best cross-validation results on the benchmark dataset. The following GNN types were considered: Graph Attention Network that fixes the static attention problem (GAT) (Brody et al., 2021), Graph Convolutional Network (GCN) (Kipf and Welling, 2016), Graph Isomorphism Network (GIN) (Xu et al., 2018), Graph Isomorphism Network with incorporated edge features (GINE) (Hu et al., 2019), and GCN for learning molecular fingerprints (GMF) (Duvenaud et al., 2015).

Besides GNN types, the subjects to tuning included: the configuration of FC layers after 1D input data, GNN pooling output and final Dense Neural Network; dropout rates; activation function for GNN layers; usage of batch normalisation; graph pooling type or combination of types.

The 3DProtDTA model was built with an open-source machine learning framework PyTorch (Paszke et al., 2019) and the GNNs were implemented using PyTorch Geometric (Fey and Lenssen, 2019).

### Comparison with existing techniques

We compared the results of our approach to different classical machine learning-based and deep learning-based methods, which are considered to be state-of-art at the time of writing.

Similarity-based approaches KronRLS (Pahikkala et al., 2015) and SimBoost (He et al., 2017) used a similarity matrix computed using Pubchem structure clustering server (Pubchem Sim, http://pubchem.ncbi.nlm.nih.gov) to represent ligands and the protein similarity matrix constructed with help of Smith-Waterman algorithm to represent targets (Smith and Waterman, 1981). KronRLS uses the regularised least-square model while SimBoost is the gradient boosting machine-based method.

The DeepDTA method (Öztürk et al., 2018) used 1D CNNs to process protein sequences and SMILES of the ligands. The GANsDTA (Zhao et al., 2020) proposed a semi-supervised GANs-based method to predict binding affinity using target sequences and ligand SMILES. The same initial protein and ligand representations were used in the DeepCDA (Abbasi et al., 2020) method, where authors applied encoding by CNN and LSTM blocks. The GraphDTA (Nguyen et al., 2021) authors proposed GNNs to process ligand graphs, while proteins were still encoded by CNN applied to sequences.

### Evaluation metrics

We selected 3 evaluation metrics used by most authors of the baseline approaches.

The mean squared error (MSE):

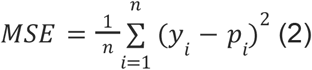

where *n* is the number of samples, *y_i_* is the observed value, and *p_i_* is the predicted value.

The concordance Index (CI) (Gönen and Heller, 2005):

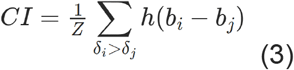

where *b_i_* is the prediction for the larger affinity *δ_i_, b_j_* is the prediction value for the smaller affinity *δ_j_*, Z is a normalisation constant, h(x) is the step function:

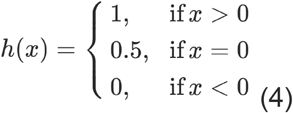

The r_m_^2^ index (Roy et al., 2013):

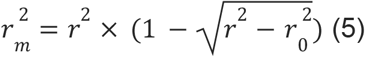

where *r^2^* and 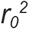 are the squared correlation coefficients with and without intercept respectively.

### Experimental setup

To assess our approach properly, we used the same train-test and cross-validation data split as in the baseline approaches. Specifically, KIBA (Tang et al., 2014) and Davis (Davis et al., 2011) datasets were divided into six equal parts in which one part is selected as the independent test set. The remaining parts of the dataset were used to determine the hyper-parameters via 5-fold cross-validation. All 5-fold training sets were used for model training. Subsequently, 5 trained models were applied to predict test set affinity. Finally, an average for each metric was calculated and compared to the baseline approaches. Train and test folds of the datasets were obtained from DeepDTA (Öztürk et al., 2018) GitHub repository.

We tuned 3DProtDTA to choose the best GNN type or GNN types combination; the number of multi-head-attentions / size of output sample (number of output node features) in GNNs; usage of activation function after a GNN layer and type of the function (ReLU, Leaky ReLU, Sigmoid); the configuration of FC layers; dropout rates; usage of batch normalisation; graph pooling type/types.

The tuning was performed with help of hyperparameter optimization software Optuna v. 3.0.3 (Akiba et al., 2019) using the Tree-structured Parzen Estimator algorithm. We trained the model for 700 epochs and used a batch size of 32, the Adam optimiser with a learning rate of 0.0001, and the Mean Squared Error Loss function.

After the tuning by the Tree-structured Parzen Estimator algorithm, we ran a range of manual cross-validation experiments keeping all the components in the best tuned architecture fixed except:

- The type of GNN architecture for a protein;
- The type of GNN architecture for a ligand;
- The type of graph pooling for both protein and ligand;
- Usage of ligand graph or Morgan fingerprint as the only ligand features.

These experiments were performed in order to assess in depth the impact of GNN architecture, graph pooling and ligand features. While changing the type of GNN model, we kept the size of output (or the number of attention heads in the case of GAT) equal or as close as possible to the tuned parameters.

## Results and discussion

### Benchmarking

The best model architectures and tuned hyperparameters for the benchmark are available in Supplementary Materials (Figure S1). All the data for model training is freely available in the GitHub repository (https://github.com/vtarasv/3d-prot-dta.git).

Tables 5 and 6 compare obtained model performance metrics to the baseline approaches for Davis and KIBA datasets respectively. The best scores are shown in bold.

**Table 5.** The average MSE, CI, and r_m_^2^ scores of the test set trained on five different training sets for the Davis dataset.

| Approach | MSE | CI | $r_m^2$ |
| --- | --- | --- | --- |
| KronRLS | 0.379 | 0.871 | 0.407 |
| SimBoost | 0.282 | 0.872 | 0.644 |
| DeepDTA | 0.261 | 0.878 | 0.63 |
| GANsDTA | 0.276 | 0.881 | 0.653 |
| DeepCDA | 0.248 | 0.891 | 0.649 |
| GraphDTA | 0.229 | 0.893 | 0.63 |
| 3DProtDTA | **0.184** | **0.917** | **0.722** |

**Table 6.**
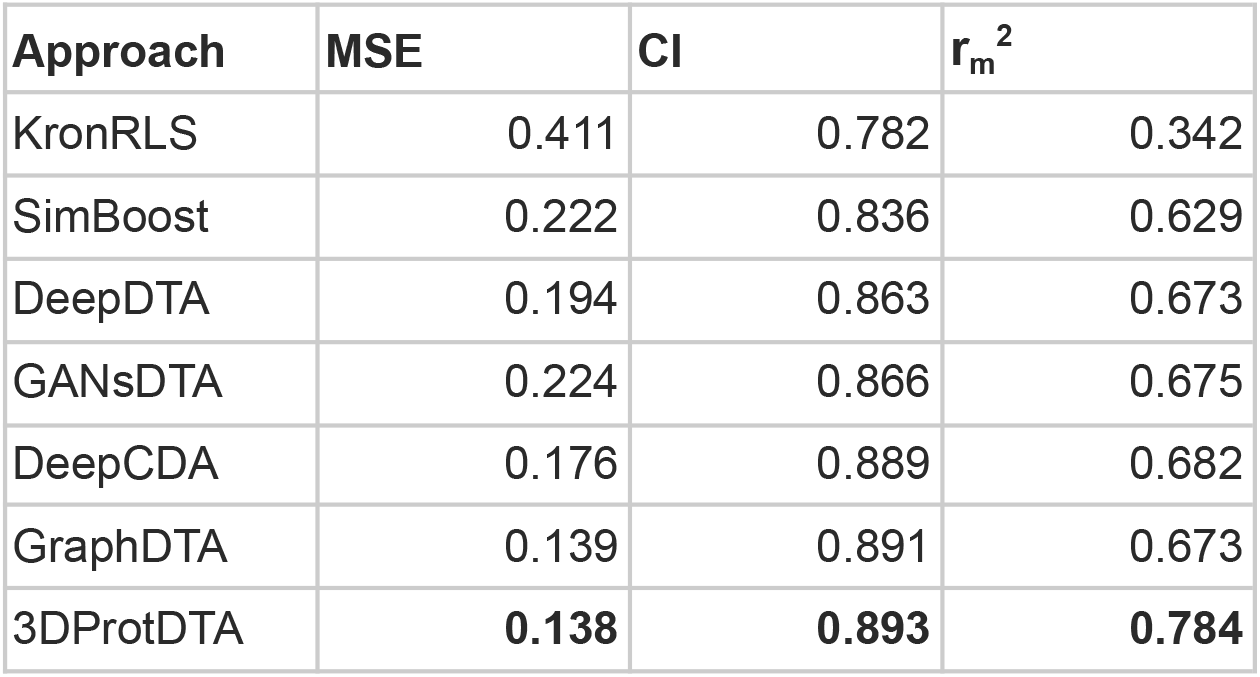
The average MSE, CI, and r_m_^2^ scores of the test set trained on five different training sets for the KIBA dataset.

According to obtained results, 3DProtDTA considerably outperforms other approaches in terms of all 3 metrics on the Davis dataset. In the case of the KIBA dataset, the only metric that demonstrated significant improvement over competitors was r_m_^2^, while the two other metrics were comparable (but still superior) to the rivals.

### Performance of ligand feature types

Atom-level molecular graphs and Morgan fingerprints are widely used types of features in the development of predictive models that take small molecules as input. The result of model training on each type separately or both types together is provided in Figure 5.

**Figure 5.**
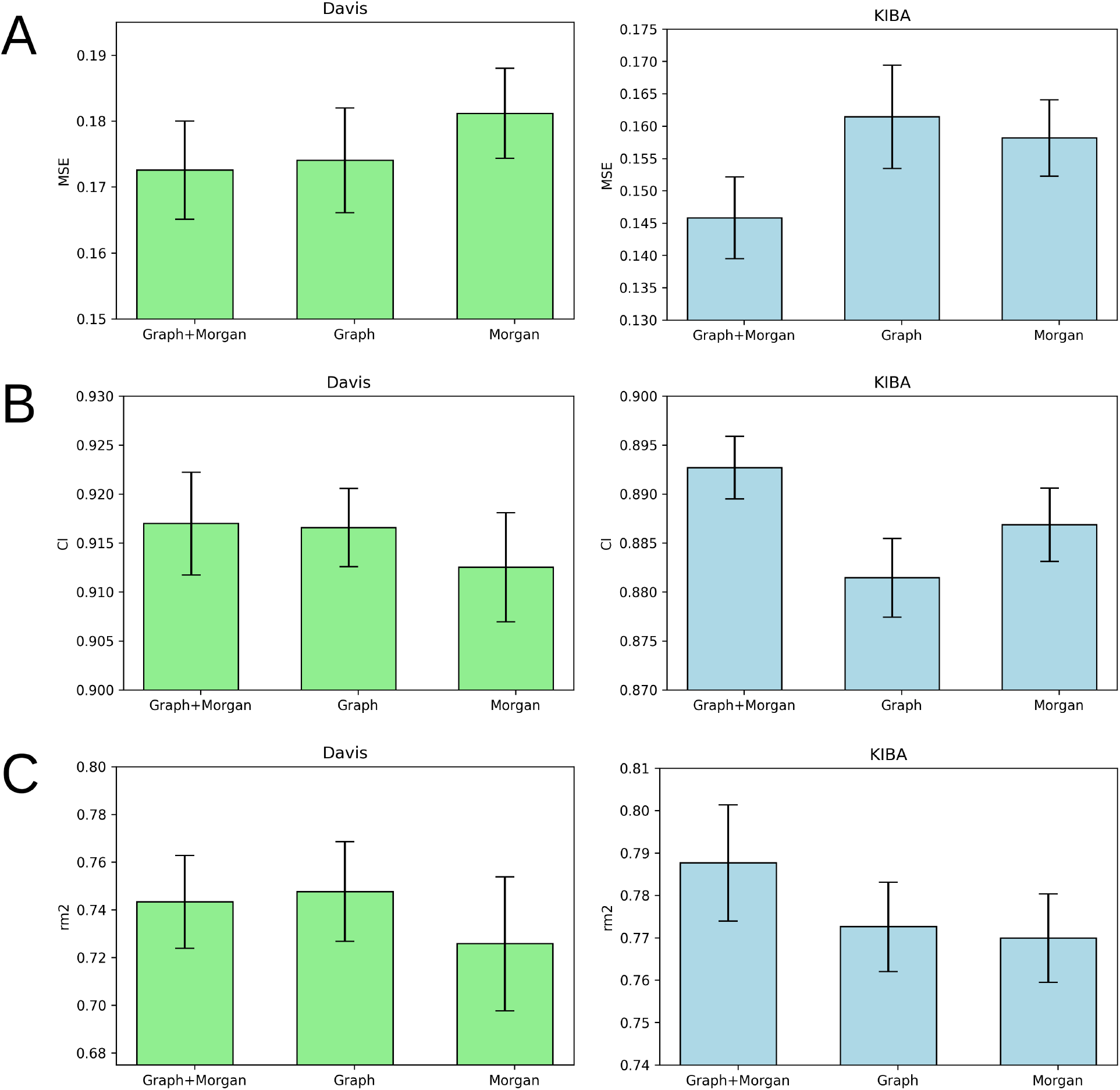
Average MSE (A), CI (B), and r_m_^2^ index (C) after 5-fold cross-validation for Davis (left) and KIBA (right) datasets. Models that use molecular graph and Morgan fingerprint (Graph+Morgan), only molecular graph (Graph) or only Morgan fingerprint (Morgan) as ligand representation are compared. Error bars represent standard deviation.

The results demonstrate the nearly equal performance of molecular graph only and combined representation of ligands for the Davis dataset. Nevertheless, the combined representation is clearly superior to others in terms of the KIBA dataset. The model trained on the molecular graph and Morgan fingerprint together provided the best average MSE, CI, and r_m_^2^ index of both benchmark datasets (Table S5). Therefore, we can suggest that these two ligand feature types contain some non-overlapping information useful for DTA prediction.

### Performance of graph pooling methods

Graph pooling is a crucial step used to generate the same-length 1D latent representation of data processed by GNN for subsequent processing by FC layers. There are three common types of pooling methods including mean pooling, max pooling and add (sum) pooling. Figure 6 provides a comparison of these pooling approaches.

**Figure 6.**
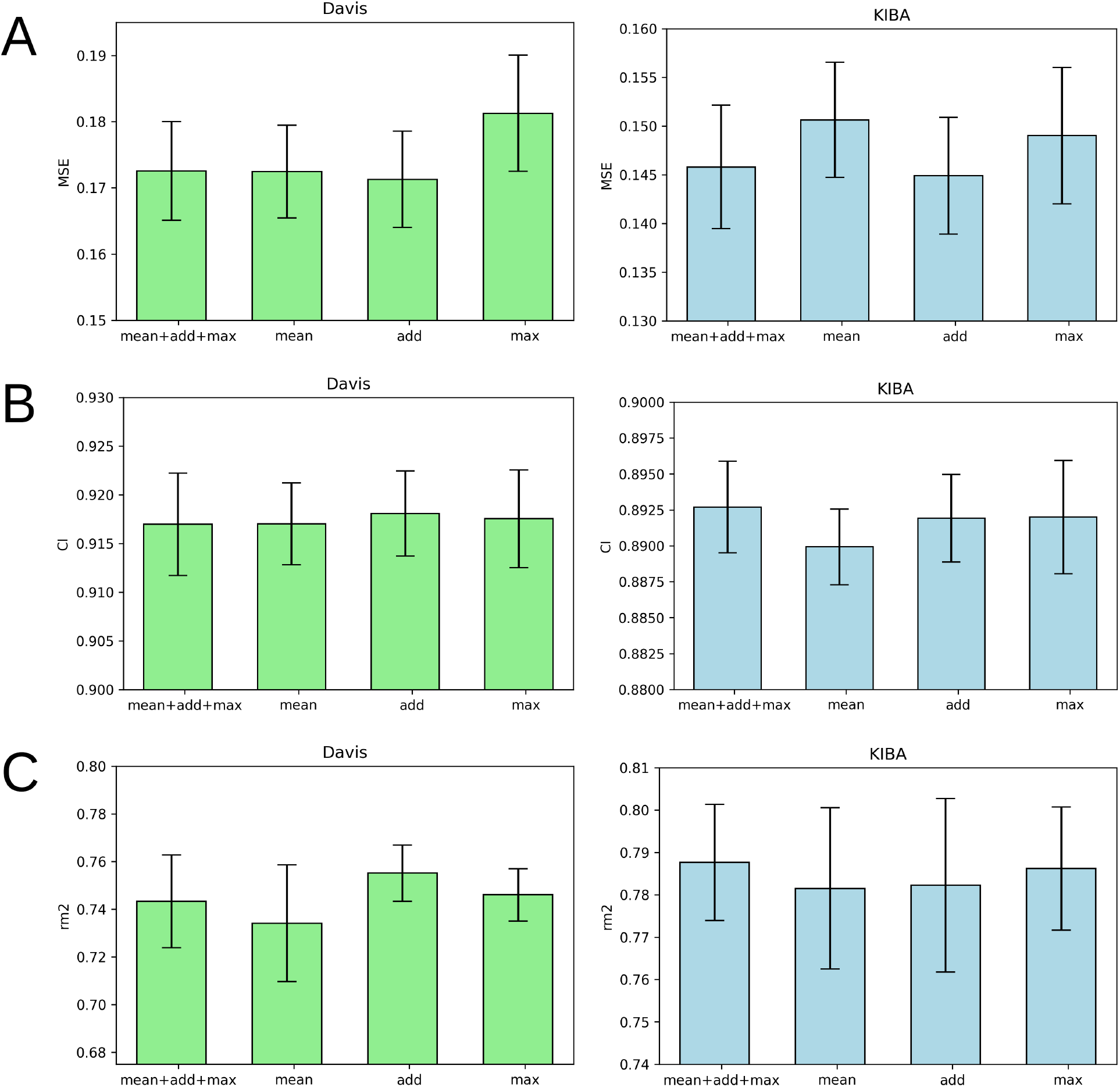
Average MSE (A), CI (B), and r_m_^2^ index (C) after 5-fold cross-validation for Davis (left) and KIBA (right) datasets. Models that use mean graph pooling (mean), add graph pooling (add), max graph pooling (max) or all three poolings concatenated (mean+add+max) are compared. Error bars represent standard deviation.

Figure 6 demonstrates that add pooling is the best choice for the Davis dataset, while add pooling is comparable to the combined approach (mean+add+max poolings concatenated) in the KIBA dataset cross-validation results. The add pooling provides superior average MSE, CI, and r_m_^2^ index of both benchmark datasets (Table S5).

### Performance of ligand GNN types

Figure 7 illustrates that there is no obvious leader or outsider as the GNN for ligand graph processing. The average of the two benchmark datasets is very close for all the GNN types considering any of the three evaluation metrics (Table S5). It is important to highlight that the only GNN model that used edge features during the training was GINE. Thus, the specific characteristic of the edge (like covalent bond type) doesn’t play a crucial role in affinity prediction.

**Figure 7.**
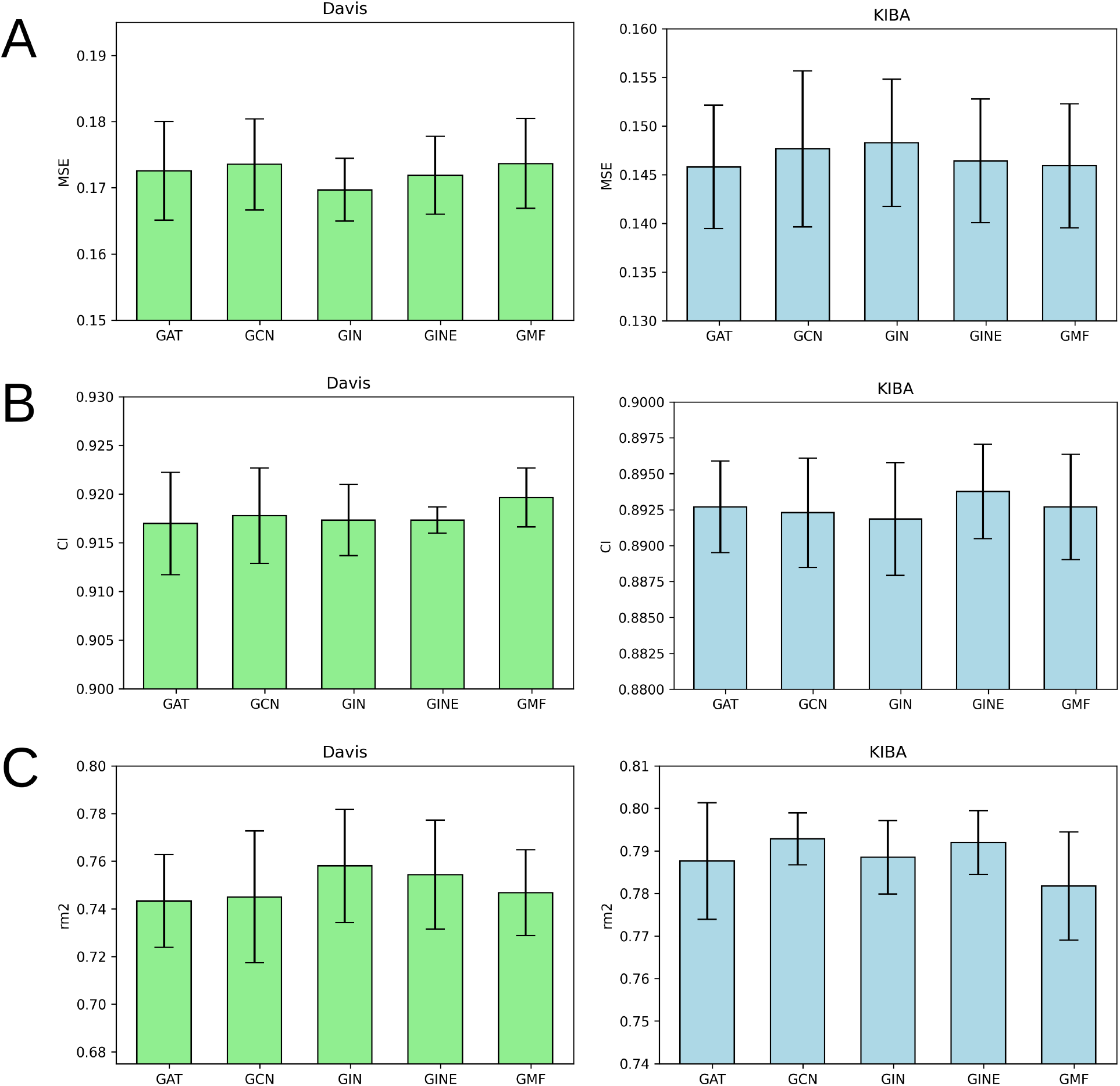
Average MSE (A), CI (B), and rm2 index (C) after 5-fold cross-validation for Davis (left) and KIBA (right) datasets. Models that use GAT, GCN, GIN, GINE or GMF to process ligand atom-level graphs are compared. Error bars represent standard deviation.

### Performance of protein GNN types

According to Figure 8, it is hard to extract a particular GNN model that works the best as a protein graph encoder. The GAT, GCN, GIN and GINE provide very similar average results. The GMF model, however, is noticeably worse (Table S5). Analogously to the performance of different GNNs on ligand graphs, the only GNN considering edge features of the protein graph GINE didn’t show considerable metrics improvement relative to other GNN types. This suggests that covalent and non-covalent interactions of amino acid residues alone provide enough information for DTA prediction.

**Figure 8.**
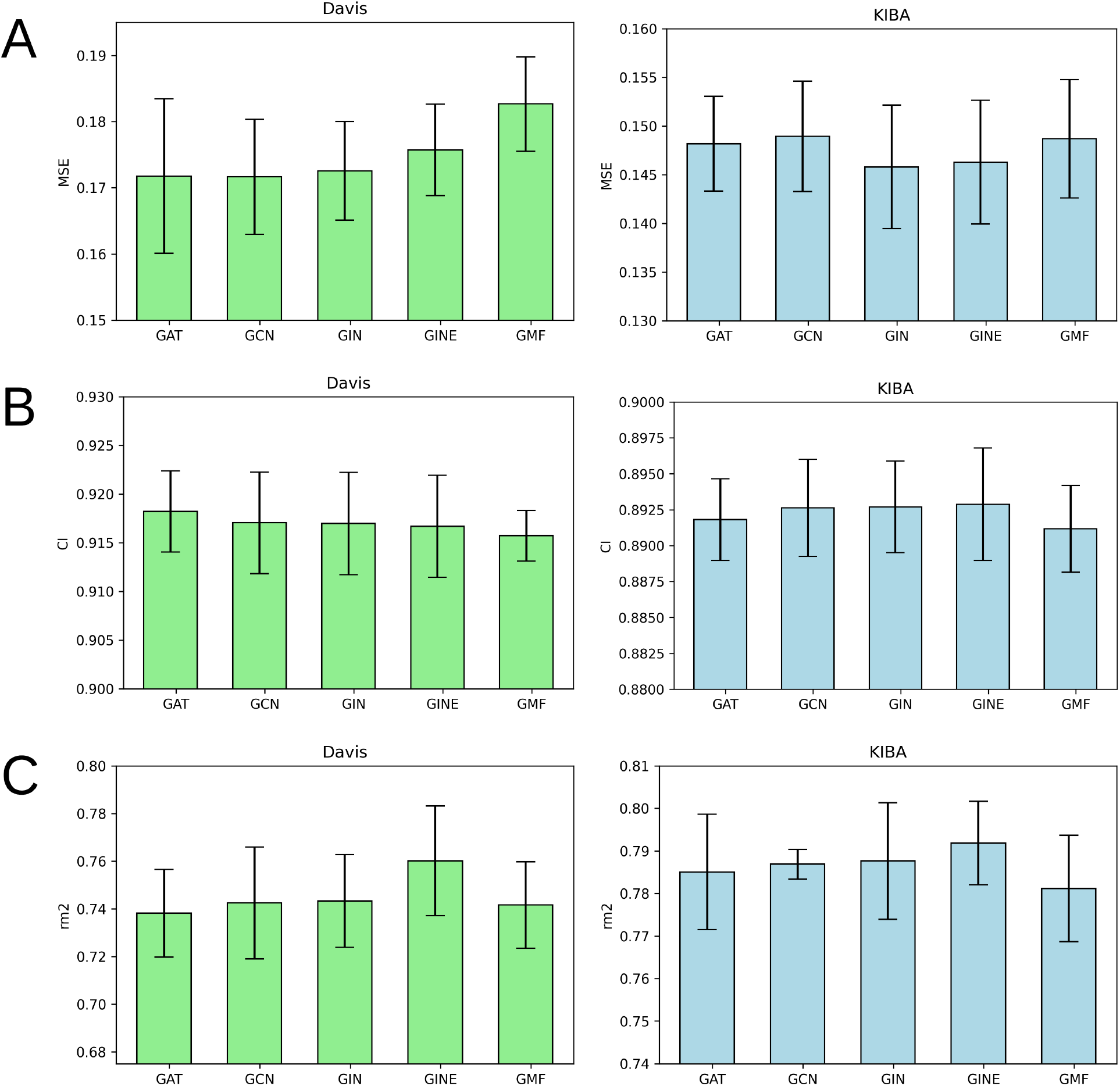
Average MSE (A), CI (B), and rm2 index (C) after 5-fold cross-validation for Davis (left) and KIBA (right) datasets. Models that use GAT, GCN, GIN, GINE or GMF to process protein residue-level graphs are compared. Error bars represent standard deviation.

### Limitations and perspectives

Although the usage of protein structures from the AlphaFold database provides data uniformity, it has some drawbacks. Particularly, this approach introduces uncertainty if multiple experimental structures are resolved and used as templates. In such cases, the AlphaFold could potentially produce a blend between several alternative protein conformations, which is not functionally relevant. An obvious improvement would be the usage of all available experimentally determined protein structures along with the AlphaFold predictions.

Another straightforward improvement is the usage of atomic-level protein graphs instead of residue-level ones, and the usage of more comprehensive node and edge features. Particularly, the B-factors and other measures of protein flexibility, such as cross-correlation matrices of motions, could be used.

## Conclusions

In this work we developed a new deep learning model for assessing drug-protein affinities called 3DProtDTA. The distinctive feature of this model is graph-based representation of both protein and the ligands, which retain a significant amount of information about their connectivity and spatial arrangement without introducing excessive computational burden. The features for model training were created using the AlphaFold database of predicted protein structures which allows for covering all proteins in two common benchmark datasets. We tuned a wide range of GNN-based model architectures and their combinations to achieve the best model performance. The 3DProtDTA outperforms its competitors on common benchmarking datasets and has a potential for further improvement.

## Author contributions

SS and AN designed the study and supervised model development and testing. SY coordinated the work, provided scripts for Pteros molecular modelling library and participated in results interpretation. TV developed the modules for features generation, model tuning and testing. TV and RS researched existing methods for drug-target affinity prediction. DN and IK participated in protein preprocessing before feature generation. LP, ZO, and RZ participated in consultations about the best practices and strategies for model tuning and efficient training. PH, IK, and VV performed and supervised the tuning and testing process. The manuscript was written by TV and SY.

## Code availability

https://github.com/vtarasv/3d-prot-dta.git

## Supplementary Materials

**Figure S1.**
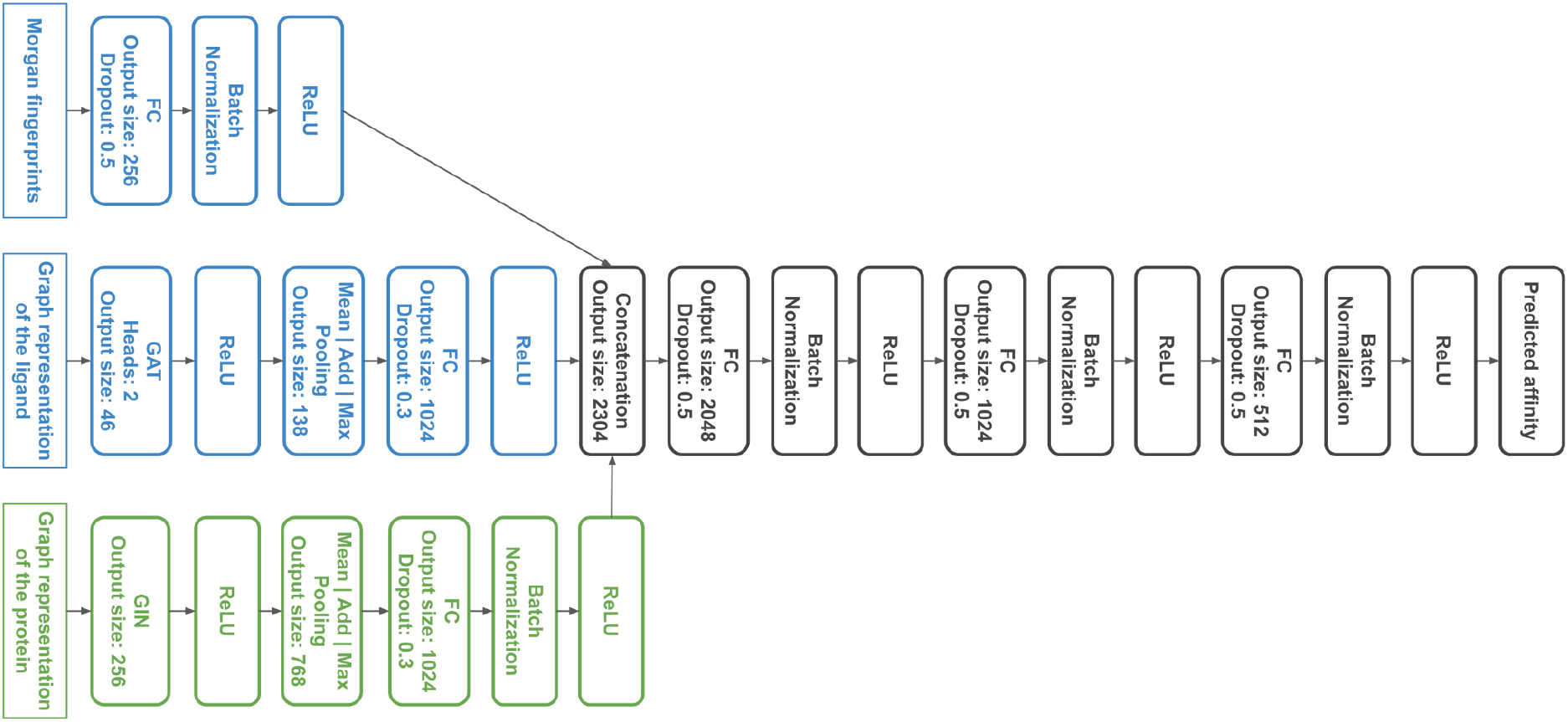
3DProtDTA model architecture tuned by Tree-structured Parzen Estimator algorithm.

**Table S1.** Detailed 5-fold cross-validation results for the Davis dataset.

| dataset | experiment | case | fold | mse | ci | rm2 |
| --- | --- | --- | --- | --- | --- | --- |
| Davis | ligand features | Graph+Morgan | 1 | 0.178 | 0.910 | 0.759 |
| Davis | ligand features | Graph+Morgan | 2 | 0.166 | 0.920 | 0.769 |
| Davis | ligand features | Graph+Morgan | 3 | 0.172 | 0.918 | 0.725 |
| Davis | ligand features | Graph+Morgan | 4 | 0.165 | 0.923 | 0.730 |
| Davis | ligand features | Graph+Morgan | 5 | 0.182 | 0.913 | 0.733 |
| Davis | ligand features | Graph | 1 | 0.177 | 0.912 | 0.765 |
| Davis | ligand features | Graph | 2 | 0.173 | 0.918 | 0.759 |
| Davis | ligand features | Graph | 3 | 0.177 | 0.920 | 0.713 |
| Davis | ligand features | Graph | 4 | 0.161 | 0.921 | 0.758 |
| Davis | ligand features | Graph | 5 | 0.182 | 0.912 | 0.744 |
| Davis | ligand features | Morgan | 1 | 0.189 | 0.908 | 0.744 |
| Davis | ligand features | Morgan | 2 | 0.176 | 0.907 | 0.758 |
| Davis | ligand features | Morgan | 3 | 0.181 | 0.919 | 0.684 |
| Davis | ligand features | Morgan | 4 | 0.173 | 0.918 | 0.724 |
| Davis | ligand features | Morgan | 5 | 0.186 | 0.911 | 0.719 |
| Davis | graph pooling | mean+add+max | 1 | 0.178 | 0.910 | 0.759 |
| Davis | graph pooling | mean+add+max | 2 | 0.166 | 0.920 | 0.769 |
| Davis | graph pooling | mean+add+max | 3 | 0.172 | 0.918 | 0.725 |
| Davis | graph pooling | mean+add+max | 4 | 0.165 | 0.923 | 0.730 |
| Davis | graph pooling | mean+add+max | 5 | 0.182 | 0.913 | 0.733 |
| Davis | graph pooling | mean | 1 | 0.173 | 0.912 | 0.752 |
| Davis | graph pooling | mean | 2 | 0.168 | 0.919 | 0.754 |
| Davis | graph pooling | mean | 3 | 0.177 | 0.921 | 0.695 |
| Davis | graph pooling | mean | 4 | 0.163 | 0.920 | 0.744 |
| Davis | graph pooling | mean | 5 | 0.181 | 0.914 | 0.725 |
| Davis | graph pooling | add | 1 | 0.173 | 0.913 | 0.753 |
| Davis | graph pooling | add | 2 | 0.166 | 0.921 | 0.769 |
| Davis | graph pooling | add | 3 | 0.175 | 0.918 | 0.738 |
| Davis | graph pooling | add | 4 | 0.162 | 0.924 | 0.752 |
| Davis | graph pooling | add | 5 | 0.181 | 0.915 | 0.763 |
| Davis | graph pooling | max | 1 | 0.185 | 0.912 | 0.757 |
| Davis | graph pooling | max | 2 | 0.174 | 0.925 | 0.757 |
| Davis | graph pooling | max | 3 | 0.177 | 0.917 | 0.732 |
| Davis | graph pooling | max | 4 | 0.175 | 0.920 | 0.745 |
| Davis | graph pooling | max | 5 | 0.195 | 0.914 | 0.738 |
| Davis | ligand GNN | GAT | 1 | 0.178 | 0.910 | 0.759 |
| Davis | ligand GNN | GAT | 2 | 0.166 | 0.920 | 0.769 |
| Davis | ligand GNN | GAT | 3 | 0.172 | 0.918 | 0.725 |
| Davis | ligand GNN | GAT | 4 | 0.165 | 0.923 | 0.730 |
| Davis | ligand GNN | GAT | 5 | 0.182 | 0.913 | 0.733 |
| Davis | ligand GNN | GCN | 1 | 0.172 | 0.913 | 0.775 |
| Davis | ligand GNN | GCN | 2 | 0.168 | 0.917 | 0.766 |
| Davis | ligand GNN | GCN | 3 | 0.182 | 0.922 | 0.704 |
| Davis | ligand GNN | GCN | 4 | 0.166 | 0.924 | 0.738 |
| Davis | ligand GNN | GCN | 5 | 0.179 | 0.913 | 0.743 |
| Davis | ligand GNN | GIN | 1 | 0.170 | 0.915 | 0.775 |
| Davis | ligand GNN | GIN | 2 | 0.163 | 0.916 | 0.778 |
| Davis | ligand GNN | GIN | 3 | 0.173 | 0.918 | 0.724 |
| Davis | ligand GNN | GIN | 4 | 0.167 | 0.923 | 0.771 |
| Davis | ligand GNN | GIN | 5 | 0.175 | 0.914 | 0.742 |
| Davis | ligand GNN | GINE | 1 | 0.171 | 0.917 | 0.768 |
| Davis | ligand GNN | GINE | 2 | 0.165 | 0.917 | 0.760 |
| Davis | ligand GNN | GINE | 3 | 0.176 | 0.918 | 0.714 |
| Davis | ligand GNN | GINE | 4 | 0.168 | 0.919 | 0.764 |
| Davis | ligand GNN | GINE | 5 | 0.179 | 0.915 | 0.765 |
| Davis | ligand GNN | GMF | 1 | 0.174 | 0.917 | 0.765 |
| Davis | ligand GNN | GMF | 2 | 0.173 | 0.922 | 0.736 |
| Davis | ligand GNN | GMF | 3 | 0.178 | 0.922 | 0.722 |
| Davis | ligand GNN | GMF | 4 | 0.163 | 0.921 | 0.750 |
| Davis | ligand GNN | GMF | 5 | 0.181 | 0.916 | 0.762 |
| Davis | protein GNN | GAT | 1 | 0.170 | 0.914 | 0.748 |
| Davis | protein GNN | GAT | 2 | 0.166 | 0.920 | 0.742 |
| Davis | protein GNN | GAT | 3 | 0.177 | 0.923 | 0.705 |
| Davis | protein GNN | GAT | 4 | 0.158 | 0.921 | 0.747 |
| Davis | protein GNN | GAT | 5 | 0.188 | 0.913 | 0.747 |
| Davis | protein GNN | GCN | 1 | 0.173 | 0.912 | 0.752 |
| Davis | protein GNN | GCN | 2 | 0.166 | 0.915 | 0.774 |
| Davis | protein GNN | GCN | 3 | 0.172 | 0.921 | 0.710 |
| Davis | protein GNN | GCN | 4 | 0.162 | 0.924 | 0.739 |
| Davis | protein GNN | GCN | 5 | 0.185 | 0.914 | 0.737 |
| Davis | protein GNN | GIN | 1 | 0.178 | 0.910 | 0.759 |
| Davis | protein GNN | GIN | 2 | 0.166 | 0.920 | 0.769 |
| Davis | protein GNN | GIN | 3 | 0.172 | 0.918 | 0.725 |
| Davis | protein GNN | GIN | 4 | 0.165 | 0.923 | 0.730 |
| Davis | protein GNN | GIN | 5 | 0.182 | 0.913 | 0.733 |
| Davis | protein GNN | GINE | 1 | 0.178 | 0.913 | 0.790 |
| Davis | protein GNN | GINE | 2 | 0.169 | 0.917 | 0.770 |
| Davis | protein GNN | GINE | 3 | 0.178 | 0.917 | 0.727 |
| Davis | protein GNN | GINE | 4 | 0.168 | 0.925 | 0.762 |
| Davis | protein GNN | GINE | 5 | 0.185 | 0.911 | 0.752 |
| Davis | protein GNN | GMF | 1 | 0.186 | 0.915 | 0.736 |
| Davis | protein GNN | GMF | 2 | 0.178 | 0.918 | 0.760 |
| Davis | protein GNN | GMF | 3 | 0.182 | 0.916 | 0.718 |
| Davis | protein GNN | GMF | 4 | 0.174 | 0.918 | 0.760 |
| Davis | protein GNN | GMF | 5 | 0.193 | 0.911 | 0.734 |

**Table S2.** Detailed 5-fold cross-validation results for the KIBA dataset.

| dataset | experiment | case | fold | mse | ci | rm2 |
| --- | --- | --- | --- | --- | --- | --- |
| KIBA | ligand features | Graph+Morgan | 1 | 0.137 | 0.897 | 0.805 |
| KIBA | ligand features | Graph+Morgan | 2 | 0.147 | 0.892 | 0.789 |
| KIBA | ligand features | Graph+Morgan | 3 | 0.145 | 0.895 | 0.796 |
| KIBA | ligand features | Graph+Morgan | 4 | 0.154 | 0.890 | 0.772 |
| KIBA | ligand features | Graph+Morgan | 5 | 0.146 | 0.890 | 0.777 |
| KIBA | ligand features | Graph | 1 | 0.150 | 0.886 | 0.788 |
| KIBA | ligand features | Graph | 2 | 0.162 | 0.883 | 0.774 |
| KIBA | ligand features | Graph | 3 | 0.165 | 0.883 | 0.764 |
| KIBA | ligand features | Graph | 4 | 0.172 | 0.876 | 0.762 |
| KIBA | ligand features | Graph | 5 | 0.159 | 0.879 | 0.776 |
| KIBA | ligand features | Morgan | 1 | 0.151 | 0.891 | 0.783 |
| KIBA | ligand features | Morgan | 2 | 0.160 | 0.888 | 0.771 |
| KIBA | ligand features | Morgan | 3 | 0.158 | 0.888 | 0.776 |
| KIBA | ligand features | Morgan | 4 | 0.167 | 0.881 | 0.756 |
| KIBA | ligand features | Morgan | 5 | 0.156 | 0.885 | 0.764 |
| KIBA | graph pooling | mean+add+max | 1 | 0.137 | 0.897 | 0.805 |
| KIBA | graph pooling | mean+add+max | 2 | 0.147 | 0.892 | 0.789 |
| KIBA | graph pooling | mean+add+max | 3 | 0.145 | 0.895 | 0.796 |
| KIBA | graph pooling | mean+add+max | 4 | 0.154 | 0.890 | 0.772 |
| KIBA | graph pooling | mean+add+max | 5 | 0.146 | 0.890 | 0.777 |
| KIBA | graph pooling | mean | 1 | 0.143 | 0.893 | 0.792 |
| KIBA | graph pooling | mean | 2 | 0.151 | 0.890 | 0.776 |
| KIBA | graph pooling | mean | 3 | 0.151 | 0.892 | 0.794 |
| KIBA | graph pooling | mean | 4 | 0.160 | 0.886 | 0.751 |
| KIBA | graph pooling | mean | 5 | 0.149 | 0.889 | 0.795 |
| KIBA | graph pooling | add | 1 | 0.138 | 0.895 | 0.795 |
| KIBA | graph pooling | add | 2 | 0.147 | 0.892 | 0.793 |
| KIBA | graph pooling | add | 3 | 0.143 | 0.894 | 0.778 |
| KIBA | graph pooling | add | 4 | 0.154 | 0.888 | 0.748 |
| KIBA | graph pooling | add | 5 | 0.142 | 0.891 | 0.797 |
| KIBA | graph pooling | max | 1 | 0.139 | 0.897 | 0.804 |
| KIBA | graph pooling | max | 2 | 0.154 | 0.892 | 0.767 |
| KIBA | graph pooling | max | 3 | 0.150 | 0.894 | 0.796 |
| KIBA | graph pooling | max | 4 | 0.157 | 0.886 | 0.777 |
| KIBA | graph pooling | max | 5 | 0.146 | 0.891 | 0.787 |
| KIBA | ligand GNN | GAT | 1 | 0.137 | 0.897 | 0.805 |
| KIBA | ligand GNN | GAT | 2 | 0.147 | 0.892 | 0.789 |
| KIBA | ligand GNN | GAT | 3 | 0.145 | 0.895 | 0.796 |
| KIBA | ligand GNN | GAT | 4 | 0.154 | 0.890 | 0.772 |
| KIBA | ligand GNN | GAT | 5 | 0.146 | 0.890 | 0.777 |
| KIBA | ligand GNN | GCN | 1 | 0.138 | 0.897 | 0.796 |
| KIBA | ligand GNN | GCN | 2 | 0.151 | 0.893 | 0.790 |
| KIBA | ligand GNN | GCN | 3 | 0.147 | 0.894 | 0.800 |
| KIBA | ligand GNN | GCN | 4 | 0.159 | 0.887 | 0.784 |
| KIBA | ligand GNN | GCN | 5 | 0.143 | 0.890 | 0.794 |
| KIBA | ligand GNN | GIN | 1 | 0.140 | 0.897 | 0.796 |
| KIBA | ligand GNN | GIN | 2 | 0.148 | 0.891 | 0.793 |
| KIBA | ligand GNN | GIN | 3 | 0.149 | 0.895 | 0.791 |
| KIBA | ligand GNN | GIN | 4 | 0.158 | 0.887 | 0.774 |
| KIBA | ligand GNN | GIN | 5 | 0.145 | 0.889 | 0.788 |
| KIBA | ligand GNN | GINE | 1 | 0.138 | 0.897 | 0.801 |
| KIBA | ligand GNN | GINE | 2 | 0.146 | 0.895 | 0.786 |
| KIBA | ligand GNN | GINE | 3 | 0.147 | 0.895 | 0.792 |
| KIBA | ligand GNN | GINE | 4 | 0.156 | 0.889 | 0.783 |
| KIBA | ligand GNN | GINE | 5 | 0.145 | 0.891 | 0.797 |
| KIBA | ligand GNN | GMF | 1 | 0.138 | 0.897 | 0.784 |
| KIBA | ligand GNN | GMF | 2 | 0.149 | 0.892 | 0.764 |
| KIBA | ligand GNN | GMF | 3 | 0.145 | 0.896 | 0.796 |
| KIBA | ligand GNN | GMF | 4 | 0.155 | 0.887 | 0.774 |
| KIBA | ligand GNN | GMF | 5 | 0.143 | 0.892 | 0.790 |
| KIBA | protein GNN | GAT | 1 | 0.144 | 0.896 | 0.794 |
| KIBA | protein GNN | GAT | 2 | 0.152 | 0.892 | 0.772 |
| KIBA | protein GNN | GAT | 3 | 0.147 | 0.892 | 0.792 |
| KIBA | protein GNN | GAT | 4 | 0.154 | 0.888 | 0.769 |
| KIBA | protein GNN | GAT | 5 | 0.144 | 0.891 | 0.798 |
| KIBA | protein GNN | GCN | 1 | 0.140 | 0.897 | 0.790 |
| KIBA | protein GNN | GCN | 2 | 0.150 | 0.893 | 0.783 |
| KIBA | protein GNN | GCN | 3 | 0.150 | 0.893 | 0.789 |
| KIBA | protein GNN | GCN | 4 | 0.156 | 0.888 | 0.783 |
| KIBA | protein GNN | GCN | 5 | 0.148 | 0.891 | 0.790 |
| KIBA | protein GNN | GIN | 1 | 0.137 | 0.897 | 0.805 |
| KIBA | protein GNN | GIN | 2 | 0.147 | 0.892 | 0.789 |
| KIBA | protein GNN | GIN | 3 | 0.145 | 0.895 | 0.796 |
| KIBA | protein GNN | GIN | 4 | 0.154 | 0.890 | 0.772 |
| KIBA | protein GNN | GIN | 5 | 0.146 | 0.890 | 0.777 |
| KIBA | protein GNN | GINE | 1 | 0.138 | 0.898 | 0.799 |
| KIBA | protein GNN | GINE | 2 | 0.150 | 0.893 | 0.789 |
| KIBA | protein GNN | GINE | 3 | 0.145 | 0.894 | 0.797 |
| KIBA | protein GNN | GINE | 4 | 0.155 | 0.888 | 0.776 |
| KIBA | protein GNN | GINE | 5 | 0.143 | 0.891 | 0.798 |
| KIBA | protein GNN | GMF | 1 | 0.140 | 0.896 | 0.794 |
| KIBA | protein GNN | GMF | 2 | 0.150 | 0.892 | 0.780 |
| KIBA | protein GNN | GMF | 3 | 0.149 | 0.891 | 0.782 |
| KIBA | protein GNN | GMF | 4 | 0.157 | 0.887 | 0.761 |
| KIBA | protein GNN | GMF | 5 | 0.146 | 0.890 | 0.790 |

**Table S3.**
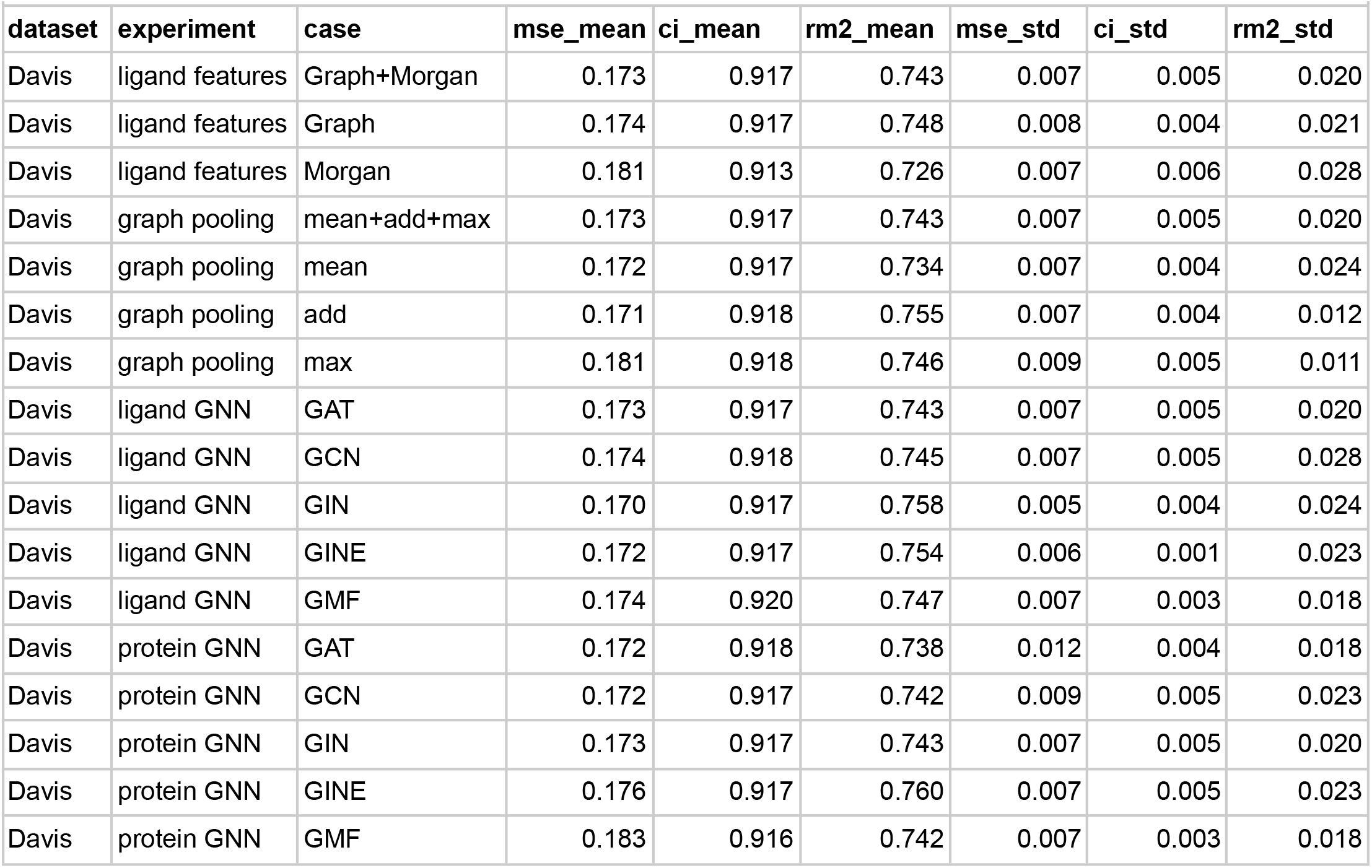
5-fold mean and standard deviation for each experiment, Davis dataset.

| dataset | experiment | case | mse_mean | ci_mean | rm2_mean | mse_std | ci_std | rm2_std |
| --- | --- | --- | --- | --- | --- | --- | --- | --- |
| Davis | ligand features | Graph+Morgan | 0.173 | 0.917 | 0.743 | 0.007 | 0.005 | 0.020 |
| Davis | ligand features | Graph | 0.174 | 0.917 | 0.748 | 0.008 | 0.004 | 0.021 |
| Davis | ligand features | Morgan | 0.181 | 0.913 | 0.726 | 0.007 | 0.006 | 0.028 |
| Davis | graph pooling | mean+add+max | 0.173 | 0.917 | 0.743 | 0.007 | 0.005 | 0.020 |
| Davis | graph pooling | mean | 0.172 | 0.917 | 0.734 | 0.007 | 0.004 | 0.024 |
| Davis | graph pooling | add | 0.171 | 0.918 | 0.755 | 0.007 | 0.004 | 0.012 |
| Davis | graph pooling | max | 0.181 | 0.918 | 0.746 | 0.009 | 0.005 | 0.011 |
| Davis | ligand GNN | GAT | 0.173 | 0.917 | 0.743 | 0.007 | 0.005 | 0.020 |
| Davis | ligand GNN | GCN | 0.174 | 0.918 | 0.745 | 0.007 | 0.005 | 0.028 |
| Davis | ligand GNN | GIN | 0.170 | 0.917 | 0.758 | 0.005 | 0.004 | 0.024 |
| Davis | ligand GNN | GINE | 0.172 | 0.917 | 0.754 | 0.006 | 0.001 | 0.023 |
| Davis | ligand GNN | GMF | 0.174 | 0.920 | 0.747 | 0.007 | 0.003 | 0.018 |
| Davis | protein GNN | GAT | 0.172 | 0.918 | 0.738 | 0.012 | 0.004 | 0.018 |
| Davis | protein GNN | GCN | 0.172 | 0.917 | 0.742 | 0.009 | 0.005 | 0.023 |
| Davis | protein GNN | GIN | 0.173 | 0.917 | 0.743 | 0.007 | 0.005 | 0.020 |
| Davis | protein GNN | GINE | 0.176 | 0.917 | 0.760 | 0.007 | 0.005 | 0.023 |
| Davis | protein GNN | GMF | 0.183 | 0.916 | 0.742 | 0.007 | 0.003 | 0.018 |

**Table S4.** 5-fold mean and standard deviation for each experiment, KIBA dataset.

| dataset | experiment | case | mse_mean | ci_mean | rm2_mean | mse_std | ci_std | rm2_std |
| --- | --- | --- | --- | --- | --- | --- | --- | --- |
| KIBA | ligand features | Graph+Morgan | 0.146 | 0.893 | 0.788 | 0.006 | 0.003 | 0.014 |
| KIBA | ligand features | Graph | 0.161 | 0.881 | 0.773 | 0.008 | 0.004 | 0.011 |
| KIBA | ligand features | Morgan | 0.158 | 0.887 | 0.770 | 0.006 | 0.004 | 0.010 |
| KIBA | graph pooling | mean+add+max | 0.146 | 0.893 | 0.788 | 0.006 | 0.003 | 0.014 |
| KIBA | graph pooling | mean | 0.151 | 0.890 | 0.781 | 0.006 | 0.003 | 0.019 |
| KIBA | graph pooling | add | 0.145 | 0.892 | 0.782 | 0.006 | 0.003 | 0.020 |
| KIBA | graph pooling | max | 0.149 | 0.892 | 0.786 | 0.007 | 0.004 | 0.015 |
| KIBA | ligand GNN | GAT | 0.146 | 0.893 | 0.788 | 0.006 | 0.003 | 0.014 |
| KIBA | ligand GNN | GCN | 0.148 | 0.892 | 0.793 | 0.008 | 0.004 | 0.006 |
| KIBA | ligand GNN | GIN | 0.148 | 0.892 | 0.789 | 0.007 | 0.004 | 0.009 |
| KIBA | ligand GNN | GINE | 0.146 | 0.894 | 0.792 | 0.006 | 0.003 | 0.007 |
| KIBA | ligand GNN | GMF | 0.146 | 0.893 | 0.782 | 0.006 | 0.004 | 0.013 |
| KIBA | protein GNN | GAT | 0.148 | 0.892 | 0.785 | 0.005 | 0.003 | 0.014 |
| KIBA | protein GNN | GCN | 0.149 | 0.893 | 0.787 | 0.006 | 0.003 | 0.004 |
| KIBA | protein GNN | GIN | 0.146 | 0.893 | 0.788 | 0.006 | 0.003 | 0.014 |
| KIBA | protein GNN | GINE | 0.146 | 0.893 | 0.792 | 0.006 | 0.004 | 0.010 |
| KIBA | protein GNN | GMF | 0.149 | 0.891 | 0.781 | 0.006 | 0.003 | 0.013 |

**Table S5.** 5-fold mean and standard deviation for each experiment, the average for the KIBA and Davis datasets.

| experiment | case | mse | ci | rm2 |
| --- | --- | --- | --- | --- |
| ligand features | Graph+Morgan | 0.159 | 0.905 | 0.765 |
| ligand features | Graph | 0.168 | 0.899 | 0.760 |
| ligand features | Morgan | 0.170 | 0.900 | 0.748 |
| graph pooling | mean+add+max | 0.159 | 0.905 | 0.765 |
| graph pooling | mean | 0.162 | 0.903 | 0.758 |
| graph pooling | add | 0.158 | 0.905 | 0.769 |
| graph pooling | max | 0.165 | 0.905 | 0.766 |
| ligand GNN | GAT | 0.159 | 0.905 | 0.765 |
| ligand GNN | GCN | 0.161 | 0.905 | 0.769 |
| ligand GNN | GIN | 0.159 | 0.905 | 0.773 |
| ligand GNN | GINE | 0.159 | 0.906 | 0.773 |
| ligand GNN | GMF | 0.160 | 0.906 | 0.764 |
| protein GNN | GAT | 0.160 | 0.905 | 0.762 |
| protein GNN | GCN | 0.160 | 0.905 | 0.765 |
| protein GNN | GIN | 0.159 | 0.905 | 0.765 |
| protein GNN | GINE | 0.161 | 0.905 | 0.776 |
| protein GNN | GMF | 0.166 | 0.903 | 0.761 |

